# Frailty index as a predictor of all-cause and cause-specific mortality in a Swedish population-based cohort

**DOI:** 10.1101/207381

**Authors:** M Jiang, AD Foebel, R Kuja-Halkola, I Karlsson, NL Pedersen, S Hägg, J Jylhävä

## Abstract

**Background:** Frailty is a complex manifestation of aging and associated with increased risk of mortality and poor health outcomes. Younger individuals (under 65 years) typically have low levels of frailty and are less-studied in this respect. Also, the relationship between the Rockwood frailty index (FI) and cause-specific mortality in community settings is understudied.

**Methods:** We created and validated a 42-item Rockwood-based FI in The Swedish Adoption/Twin Study of Aging (n=1477; 623 men, 854 women; aged 29-95 years) and analyzed its association with all-cause and cause-specific mortality in up to 30-years of follow-up. Deaths due to cardiovascular disease (CVD), cancer, dementia and other causes were considered as competing risks.

**Results:** Our FI demonstrated construct validity as its associations with age, sex and mortality were similar to the existing literature. The FI was independently associated with increased risk for all-cause mortality in younger (<65 years; HR per increase in one deficit 1.11, 95%CI 1.07-1.17) and older (≥65 years; HR 1.07, 95%CI 1.04-1.10) women and in younger men (HR 1.05, 95%CI 1.01-1.10). In cause-specific mortality analysis, the FI was strongly predictive of CVD mortality in women (HR per increase in one deficit 1.13, 95%CI 1.09-1.17), whereas in men the risk was restricted to deaths from other causes (HR 1.07, 95%CI 1.01-1.13).

**Conclusions:** The FI showed good predictive value for all-cause mortality especially in the younger group. The FI predicted CVD mortality risk in women, whereas in men it captured vulnerability to death from various causes.

## Introduction

Frailty is among the most complex and problematic conditions in older individuals. It is a state of increased vulnerability to stressors and adverse outcomes due to a multisystem loss of homeostasis. The importance of frailty is highlighted by its consistent association with allcause mortality and adverse aging outcomes, such as institutionalization, physical limitations, disability, recurrent hospitalizations, falls and fractures (1,2). In individuals aged 65 years and older, a dose-responsive reduction in the survival probability has been observed with increasing levels of frailty (1). It has been estimated that that up to 5% of deaths among older individuals could be delayed by preventing frailty (1).

There are currently two common models of frailty: the Fried frailty phenotype (3) and the Rockwood frailty index (FI) (4). The former sees frailty as a syndrome with loss of physical function and it classifies individuals as non-frail, pre-frail and frail, whereas the latter sees frailty as a continuous risk state assessed as the number of accumulated health deficits. Although both measures are valid predictors of adverse outcomes and demonstrate overlap in the identification of frailty (5), the FI - being a continuous measure - shows more sensitivity at the lower end of the frailty continuum (6). In addition, the FI may provide better resolution in younger populations as it also predicts adverse outcomes among individuals who are classified as non-frail by the Fried phenotypic model (7). Nevertheless, despite the acknowledged need to screen all individuals aged 70 years and older for frailty (8), a consensus is still lacking as to how to best assess it in clinical settings.

Whether these frailty instruments also predict mortality among the younger individuals (under 65 years), whose levels of frailty are generally much lower and times to death are longer, is less-studied. Sex differences in this respect have also received little attention. A growing body of evidence links frailty to the development of cardiovascular disease (CVD) (9,10) and dementia (11), and the presence of frailty is also known to worsen the outcomes of these diseases as well as cancer (12). However, less is known about the association of frailty with deaths due to these causes and especially in community settings. To this end, we analyzed the predictive ability of a 42-item Rockwood-based FI, herein referred to as the FI, for all-cause mortality in a population-based cohort of individuals aged from 29 to 95 years at baseline with a 30-year mortality follow-up. We also elucidated the relationship between the FI and CVD, cancer, dementia and other-cause mortality.

## Methods

### Participants

The Swedish Adoption/Twin Study of Aging (SATSA) (13) is a longitudinal population-based cohort of same-sexed twins, that is drawn from the Swedish Twin Registry (STR) (14). SATSA was initiated in 1984 (n=2018) and included both mailed questionnaires and inperson testing waves. The present study involves both single and pair responders (n=1477; 623 men, 854 women; aged 29-95 years) who returned the second questionnaire sent out in 1987. The questionnaires assess physical and mental health status, activities of daily living, wellbeing, health-related behavior with respect to smoking, alcohol consumption and use of prescribed drugs, family and social environments and personality dimensions (as documented in (15)). The study has received ethical approval from the Regional Ethics Review Board, Stockholm.

### Assessment of frailty

We constructed the FI in SATSA based on the self-reported questionnaire data using the Rockwood deficit accumulation model according to the standard procedure (4,16). The Rockwood FI uses health deficits that can be defined as symptoms, signs, disabilities and diseases that cover a wide range of systems, associate with health status and have a prevalence of ≥1% in the study population. The selected 42 items and scoring of the deficits are described in the Supplementary Table 1. All items had ≤10% missing data points and only individuals with ≤20% missing answers across the 42 FI items were included. After that, the patterns of missingness were examined, missing data were replaced by multiple imputation (MI) and a sensitivity analysis for the imputed data was performed (Supplementary Methods). Each individual’s FI was assessed by counting the number of deficits and dividing the count by the total number of deficits considered. The validity of the FI was assessed by examining its distribution and associations with age and sex. In addition, we tested whether at each given level of frailty, men were more likely to die than women. As further validation to test for a dose-response relationship with the FI and mortality, we subdivided the FI into four categories according to Rockwood et al. (2011): relatively fit (FI≤0.03), less fit (0.03<FI≤0.10), least fit (0.10<FI≤0.21) and frail (FI>0.21) (16).

### Mortality

All-cause mortality data, including dates of death, were obtained from linkages of the STR to Swedish national registers through the personal identification number assigned to all residents. The all-cause mortality data used in this study were updated on April 30, 2017, yielding a 30-year follow-up. Cause-specific mortality data were obtained from the Causes of Death Register (CDR) where the latest update was on December 31, 2014, producing a 27-year follow-up period. The CDR records include information about the underlying and contributory causes of death for all individuals who were registered as Swedish residents in the year of their death. Causes of death are recorded using the International Classification of Diseases (ICD) codes, with ICD-7 used prior to 1969, ICD-8 between 1969 and 1986, ICD-9 from 1987 until 1996, and ICD-10 from 1997 and onwards. We considered cancer, CVD (including stroke) and dementia as the specific causes of death and other causes not falling into these categories were denoted as “other”. The ICD codes included in each cause are presented in the Supplementary Table 2. When more than one cause of death was recorded, a consensus classification was used (Supplementary Table 3) and a sensitivity analysis was performed (Supplementary methods). The causes of death classified as the “other” are shown in Supplementary Table 4.

### Covariates

Age at baseline, sex, education, smoking status and body mass index (BMI) were considered as covariates in the survival analyses. Education, smoking status and BMI were assessed from questionnaire data. Education level was classified as 1=elementary school (reference category), 2=O level, vocational school or folk high school, 3=secondary education (gymnasium or A level) and 4=university or higher. Smoking status was classified as nonsmoker (reference category), ex-smoker and current smoker. BMI was calculated as weight divided by height squared (kg/m2).

### Statistical analyses

The associations between the study variables and sex were tested using the Mann-Whitney test or χ^2^-test when appropriate. As the FI had a skewed distribution (Supplementary Figure 1), the association between log(FI) and age was tested using Pearson’s rho. Kaplan-Meier survival plots were used to assess differences in mortality rates by sex and FI categories. Age, sex, FI, BMI, education and smoking status were first tested for their association with mortality in univariate Cox regression models for the whole study population. Significant covariates were then tested in multivariate Cox models. Those covariates that remained significantly associated with all-cause mortality in the whole population were used as the final model covariates. The final models were stratified by sex and further subdivided into the younger and older using the standard definition of old age, ≥65 years, as the cut point. In all survival models, the sum of the FI deficits (of the 42 considered) was used to obtain hazard ratios (HRs) that are interpretable as per increase by one FI deficit. In other statistical tests the sum of the deficits divided by 42 was used. Violation of the proportional hazards (PH) assumption was tested by including an interaction term with time with each variable and further inspected using the Schoenfeld and scaled Schoenfeld residual plots. If violation was observed, a time-varying coefficient (HR) was produced for that covariate.

To rigorously assess the relationship between FI and cause-specific mortality, we took two different approaches: the cause-specific hazard model (CHR) using the conventional Cox regression and a cumulative incidence function (CIF) using the subdistribution hazard ratio (SHR) proposed by Fine and Fray (17) (see Supplementary methods for details). In both approaches, deaths due to cancer, CVD, dementia and other causes were considered as the competing risks (mutually exclusive failures). If a significant association was found, an additional analysis performed by adjusting for the diagnosis of the given disease at the study baseline (see Supplementary methods). In all models, clustering of the data in twin pairs was accounted for by computing cluster robust standard errors for the coefficients. P-values <0.05 were considered statistically significant. Statistical analyses were performed using Stata version 14.1 (College Station, TX: StataCorp LP) and SPSS version 24.0 (Armonk, NY: IBM Corp).

## Results

### The FI in the study sample

Characteristics of the study population are presented in Table 1. The distribution of the FI was skewed with a long right tail (Supplementary Figure 1) and the association with age was exponential in both men and women (Figure 1a). The maximum value of FI was 0.619 in women and 0.536 in men. Although the median level of frailty was relatively low in this sample (Table 1), sex differences across the age range and a higher mortality risk for men at all levels of FI were apparent (Figure 1b).

**Table 1.** Characteristics of the study population.

|  | **All (n=1477)** |  | **Men (n=623)** |  | **Women (n=854)** |  | **p\*** |
| --- | --- | --- | --- | --- | --- | --- | --- |
|  | Median (IQR) |  | Median (IQR) |  | Median (IQR) |  |  |
| Age (yr) | 63.2 | (21.0) | 62.9 | (19.4) | 63.5 | (21.8) | 0.037* |
| BMI (kg/m<sup>2</sup>) | 24.2 | (4.4) | 24.7 | (3.8) | 23.6 | (4.8) | <0.001* |
| Smoking status<sup>a</sup> |  |  |  |  |  |  |  |
| Never | 1032 | (70.7) | 397 | (64.0) | 635 | (75.6) |  |
| Previous | 87 | (6.0) | 52 | (8.4) | 35 | (4.2) |  |
| Current | 341 | (23.4) | 171 | (27.6) | 170 | (20.2) | <0.001* |
| Education |  |  |  |  |  |  |  |
| Elementary school<sup>a</sup> | 800 | (57.7) | 325 | (55.5) | 475 | (59.4) |  |
| O level, vocational school or folk high school | 383 | (27.6) | 148 | (25.3) | 235 | (29.4) |  |
| Secondary education (gymnasium or A level) | 109 | (7.8) | 62 | (10.6) | 47 | (5.9) |  |
| University or higher | 49 | (6.9) | 51 | (8.7) | 43 | (5.4) | <0.001* |
| FI | 0.077 | (0.095) | 0.071 | (0.083) | 0.083 | (0.107) | 0.007* |
| FI categorized<sup>a</sup> |  |  |  |  |  |  |  |
| Relatively fit | 288 | (19.5) | 122 | (19.6) | 166 | (19.4) |  |
| Less fit | 622 | (42.1) | 279 | (44.5) | 343 | (40.2) |  |
| Least fit | 401 | (27.2) | 175 | (28.1) | 226 | (26.5) |  |
| Frail | 166 | (11.2) | 47 | (7.5) | 119 | (13.9) | 0.002* |
| All-cause mortality |  |  |  |  |  |  |  |
| Died during follow-up<sup>a</sup> | 975 | (66.1) | 432 | (69.3) | 543 | (63.6) | 0.021* |
| Median time to death | 14.8 | (12.8) | 13.7 | (13.0) | 15.5 | (12.8) | 0.001* |
| Cancer mortality |  |  |  |  |  |  |  |
| Died during follow-up<sup>a</sup> | 232 | (15.7) | 143 | (23.0) | 89 | (10.4) | <0.001* |
| Median time to death | 12.2 | (11.4) | 11.5 | (12.7) | 13.1 | (9.6) | 0.223 |
| CVD mortality |  |  |  |  |  |  |  |
| Died during follow-up<sup>a</sup> | 347 | (23.5) | 160 | (25.7) | 187 | (21.9) | 0.090 |
| Median time to death | 13.7 | (12.3) | 13.2 | (12.7) | 14.6 | (12.0) | 0.081 |
| Dementia mortality |  |  |  |  |  |  |  |
| Died during follow-up<sup>a</sup> | 78 | (5.3) | 20 | (3.2) | 58 | (6.8) | 0.002 |
| Median time to death | 19.0 | (7.8) | 19.4 | (6.1) | 18.0 | (9.0) | 0.672 |
| Other causes of mortality |  |  |  |  |  |  |  |
| Died during follow-up<sup>a</sup> | 258 | (17.5) | 89 | (14.3) | 169 | (19.8) | 0.006 |
| Median time to death | 14.1 | (11.7) | 14.1 | (11.2) | 13.6 | (11.8) | 0.743 |

Note. n=1442 for information on BMI, n=1460 for smoking status and n=1386 for education.
The follow-up time was 30 years for all-cause mortality and 27 years for cause-specific mortality. Other causes of mortality include those not dying of cancer, CVD or dementia.
<sup>a</sup>Values are n (%)
\*p<0.05 for difference between men and women, Mann-Whitney's test for median values and $\chi^2$ -test for dichotomous variables
Abbreviations: BMI, body mass index; CVD, cardiovascular disease; FI, frailty index; IQR, interquartile range

**Figure 1.**
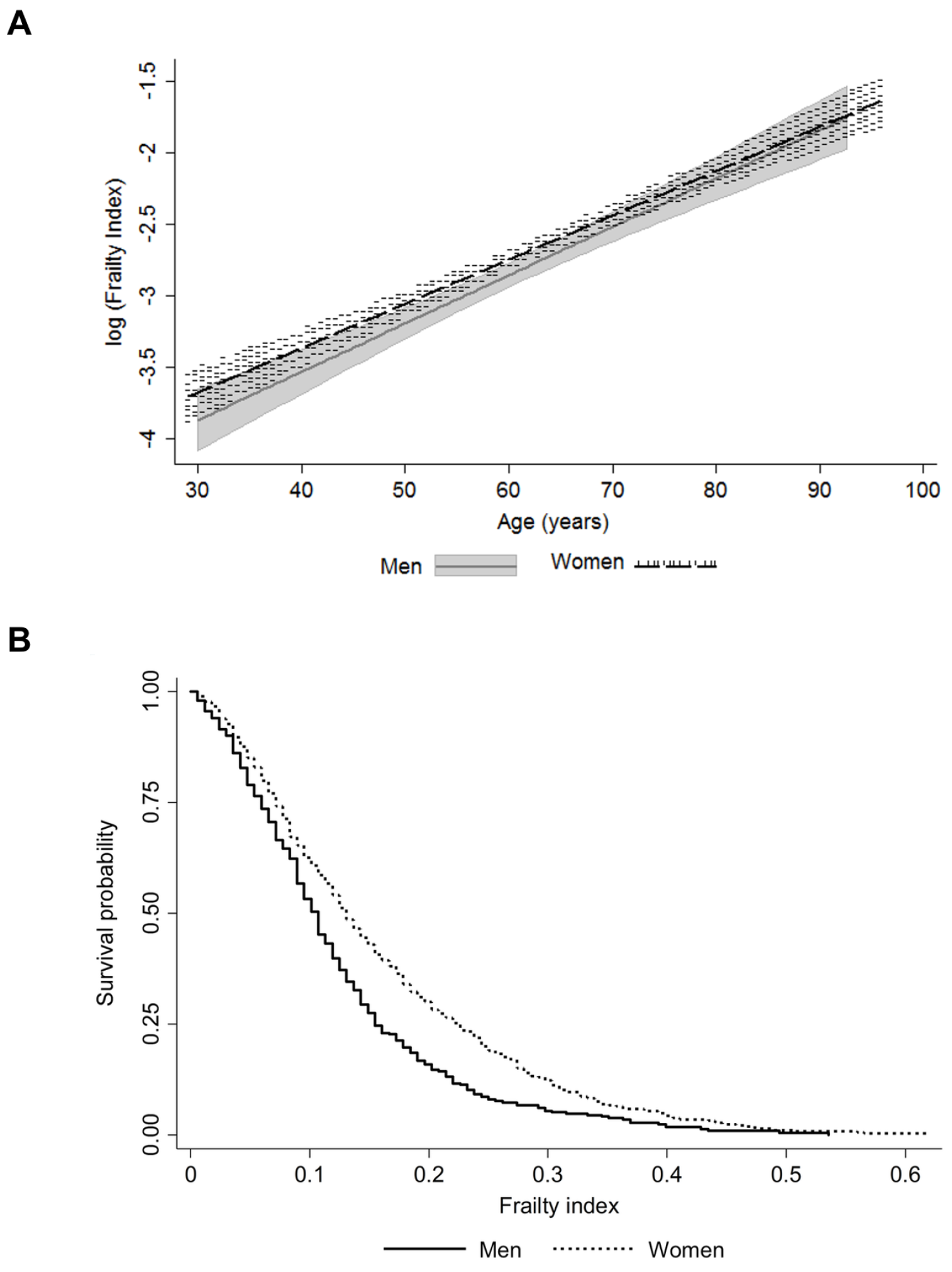
Association of the frailty index with age (a; the shaded areas around the lines represent the 95% confidence intervals for the mean) and mortality by sex (b).

### All-cause mortality

Men had higher all-cause mortality rate and a lower median time to death than women during the 30-year follow-up (Table 1). The categorized FI levels demonstrated a dose-response increase in mortality risk with increased frailty in both men and women (Supplementary Figure 2). Univariate Cox regression demonstrated that all the study variables were significantly associated with mortality in the whole population: age HR 1.13 (per year increase), 95%CI 1.12-1.14, p<0.001; sex (men as reference) HR 0.81, 95%CI 0.70-0.94, p=0.007; BMI (per unit increase) HR 1.04, 95%CI 1.02-1.06, p<0.001; education (elementary school as reference category) HR 0.68, 95%CI 0.62-0.75, p<0.001; smoking status (nonsmokers as reference) HR 0.80, 95%CI 0.74-0.87, p<0.001; and FI (sum of the deficits) HR 1.14, 95%CI 1.12-1.16, p<0.001. For BMI, only a linear association with mortality was considered, as no sign of an existence of a U-shaped relationship was observed. This was verified as the −2 log likelihood between the Cox regression model with the linear term of BMI and the model with both the linear term and the quadratic term of BMI was nonsignificant (p=0.128). Age, smoking and FI remained significantly associated with mortality in the multivariate Cox models for men and women (Table 2). When the models were further stratified by age, the associations between FI and mortality were similar in all but the older men where the association was non-significant (Table 2).

**Table 2.** Cox regression for all-cause mortality with a 30-year follow-up. Age, smoking and FIwere all assessedanalyzed in the same model.

|  | All |  |  | Young (<65 years) |  |  | Old (≥65 years) |  |  |
| --- | --- | --- | --- | --- | --- | --- | --- | --- | --- |
|  | HR | 95%CI | p | HR | 95%CI | p | HR | 95%CI | p |
| **Men** |  |  |  |  |  |  |  |  |  |
| Age | 1.12 | 1.11-1.14 | <0.001* | 1.12 | 1.09-1.15 | <0.001* | 1.11 | 1.08-1.13 | <0.001* |
| Smoking | 1.23 | 1.10-1.38 | <0.001* | 1.20 | 0.99-1.44 | 0.055 | 1.26 | 1.09-1.46 | 0.002* |
| FI | 1.04 | 1.01-1.07 | 0.008* | 1.05<sup>a</sup> | 1.01-1.10 | 0.021* | 1.03 | 0.99-1.07 | 0.097 |
| **Women** |  |  |  |  |  |  |  |  |  |
| Age | 1.13 | 1.12-1.15 | <0.001* | 1.13 | 1.10-1.16 | <0.001* | 1.14 | 1.10-1.17 | <0.001* |
| Smoking | 1.20 | 1.04-1.37 | 0.011* | 1.31 | 1.09-1.58 | 0.004* | 1.08 | 0.88-1.31 | 0.456 |
| FI | 1.08<sup>a</sup> | 1.06-1.11 | <0.001* | 1.11 | 1.07-1.17 | <0.001* | 1.07<sup>a</sup> | 1.04-1.10 | <0.001* |

Note. Numbers of deaths/individuals in each model were as follows: all men 430/620; young men 166/352; old men 264/268; all women 532/840; young women 156/452; old women 376/388.
<sup>a</sup>Proportional hazards assumption not met: a time-varying coefficient produced and presented in Supplementary Figure 3.
\*p<0.05
Abbreviations: CI, confidence interval; FI, frailty index; HR, hazard ratio

The Schoenfeld and scaled Schoenfeld residual plots with the time-varying coefficients for FI from the models where FI violated the PH assumption (younger men, all women and older women) are presented in Supplementary Figure 3. Overall, the violations were mild; no clear patterns creating the non-zero slopes were observed and the time-varying coefficients indicated 0.4-0.8% decrease in the HR for FIper year. Hence the HR for FI in these models is interpreted as a time-averaged effect.

### Cause-specific mortality

Cause-specific mortality rates and median times to death during the 27-year follow-up are presented in Table 1. The Kaplan-Meier survivor function plots for the competing risks and the median FI in each group are presented in Figure 2. The results of the age and smoking status-adjusted CHR and SHR models are presented in Table 3. In women, the CHR models demonstrated that FI predicted deaths due to cancer (HR 1.06, 95%CI 1.01-1.12), CVD (HR 1.13, 95%CI 1.09-1.17) and other causes (HR 1.07, 95%CI 1.03-1.12). In the Cox models run as a sensitivity analysis for the consensus classifications for CVD and cancer deaths, the HRs for the FI remained essentially the same (cancer mortality HR 1.06, 95%CI 1.00-1.13, p=0.05; CVD mortality HR 1.12, 95%CI 1.08-1.16), yet the former association was attenuated to borderline significance. Adjusting the CVD mortality model for CVD status at baseline did not change the results for FI (HR 1.11, 95%CI 1.06-1.16), whereas adjusting the cancer mortality model for cancer diagnosis at study baseline attenuated the FI’s association towards null (HR 1.05, 95%CI 1.00-1.11, p=0.072). In the SHR models for women, the association with CVD mortality was significant (SHR 1.07, 95%CI 1.02-1.11) and a significant inverse association was observed with dementia mortality (SHR 0.87, 95%CI 0.80-0.96). In men, the CHR models demonstrated that FI predicted only other-cause mortality (HR 1.08, 95%CI 1.02-1.14) whereas the SHR models demonstrated no significant associations.

**Table 3.** Cause-specific mortality with a 27-year follow-up. Age, smoking and FIwere all analyzed in the same model.

|  | Cause-specific hazards |  |  | Subdistribution hazards |  |  |
| --- | --- | --- | --- | --- | --- | --- |
|  | HR | 95%CI | p | SHR | 95%CI | p |
| **Men** |  |  |  |  |  |  |
| Cancer mortality |  |  |  |  |  |  |
| Age | 1.07 | 1.05-1.09 | <0.001* | 1.02 | 1.01-1.04 | 0.001* |
| Smoking | 1.19 | 0.98-1.45 | 0.087 | 1.10 | 0.91-1.34 | 0.325 |
| FI | 1.03 | 0.97-1.09 | 0.357 | 0.99 | 0.91-1.06 | 0.769 |
| CVD mortality |  |  |  |  |  |  |
| Age | 1.15 | 1.12-1.18 | <0.001* | 1.08 | 1.06-1.09 | <0.001* |
| Smoking | 1.18 | 1.04-1.37 | 0.065 | 0.97 | 0.80-1.19 | 0.777 |
| FI | 1.03 | 0.99-1.08 | 0.158 | 0.99 | 0.94-1.04 | 0.759 |
| Dementia mortality |  |  |  |  |  |  |
| Age | 1.25 | 1.17-1.33 | <0.001* | 1.05 | 1.02-1.08 | <0.001 |
| Smoking | 0.89 | 0.49-1.61 | 0.699 | 0.71 | 0.41-1.23 | 0.224 |
| FI | 1.06 | 0.91-1.23 | 0.467 | 0.97 | 0.83-1.13 | 0.672 |
| Other causes |  |  |  |  |  |  |
| Age | 1.16 | 1.12-1.19 | <0.001* | 1.06 | 1.04-1.08 | <0.001* |
| Smoking | 1.45 | 1.15-1.80 | 0.004* | 1.17 | 0.92-1.50 | 0.198 |
| FI | 1.08 | 1.02-1.14 | 0.005* | 1.04 | 0.98-1.11 | 0.178 |
| **Women** |  |  |  |  |  |  |
| Cancer mortality |  |  |  |  |  |  |
| Age | 1.08 | 1.06-1.11 | <0.001* | 1.03 | 1.02-1.05 | <0.001* |
| Smoking | 1.31 | 0.98-1.75 | 0.067 | 1.20 | 0.89-1.62 | 0.232 |
| FI | 1.06 | 1.01-1.12 | 0.031* | 1.00 | 0.95-1.05 | 0.866 |
| CVD mortality |  |  |  |  |  |  |
| Age | 1.14 | 1.11-1.16 | <0.001* | 1.06 | 1.04-1.07 | <0.001* |
| Smoking | 1.06 | 0.98-1.75 | 0.648 | 0.94 | 0.72-1.23 | 0.640 |
| FI | 1.13 | 1.09-1.17 | <0.001* | 1.07<sup>a</sup> | 1.02-1.11 | 0.001* |
| Dementia mortality |  |  |  |  |  |  |
| Age | 1.18 | 1.14-1.22 | <0.001* | 1.06 | 1.04-1.08 | <0.001* |
| Smoking | 1.23 | 0.82-1.82 | 0.315 | 0.96 | 0.64-1.44 | 0.855 |
| FI | 0.94 | 0.85-1.04 | 0.247 | 0.87 | 0.80-0.96 | 0.004* |
| Other causes |  |  |  |  |  |  |
| Age | 1.15 | 1.12-1.17 | <0.001* | 1.07 | 1.05-1.09 | <0.001* |
| Smoking | 1.37 | 1.09-1.72 | 0.007* | 1.19 | 0.95-1.49 | 0.136 |
| FI | 1.07 | 1.03-1.12 | 0.001* | 0.99 | 0.95-1.03 | 0.725 |

\*p<0.05
<sup>a</sup>Proportional hazards assumption not met: HR for FI\*time interaction=0.988
Abbreviations: CI, confidence interval; CVD, cardiovascular disease; FI, frailty index; HR, hazard ratio

**Figure 2.**
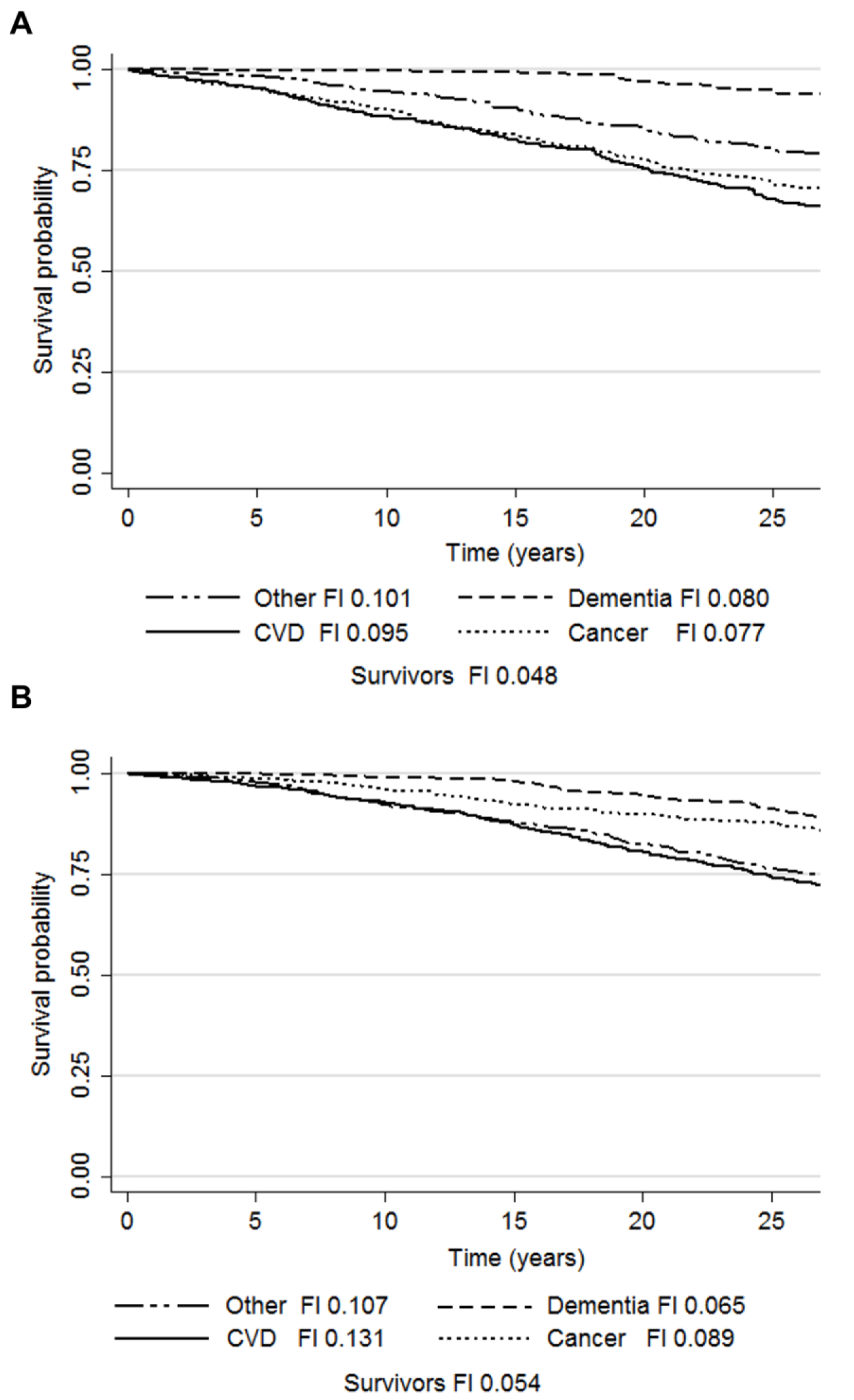
Kaplan-Meier survival probabilities and median frailty index levels according to the causes of death in men (a) and women (b). Abbreviations: CVD, cardiovascular disease; FI, frailty index.

## Discussion

In this study, we created a 42-item Rockwood FI in a population-based Swedish cohort and validated it by showing that its distribution and associations with age and sex are in agreement with those reported previously (4,6,7,16). The categorized FI also demonstrated a dose-response relationship with mortality, in keeping with the observations by Rockwood et al. (2011) (16). An increase in the FI, when treated as a continuous variable, was directly associated with all-cause mortality throughout the 30-year follow-up. The association was independent of age, smoking status, education and BMI. Stratification by sex and into younger (<65 years) and older (≥65 years) ages revealed that the association was strongest among younger women where accumulation of one deficit was associated with an 11% increase in the mortality risk. Among older women, the corresponding increase in the risk was 7%. In the younger men accumulation of one deficit was associated with a 5% increase in the mortality risk, whereas among older men the association was non-significant. The two approaches, the CHR and SHR, taken to analyze the relationships between FI and cause-specific mortality unequivocally demonstrated that the FI predicts CVD mortality in women. A suggestive association was observed for cancer mortality in women as the association was significant only in the CHR model. In men, the FI was predictive only of other-cause mortality and the finding was restricted to the CHR model.

Our results add to the understanding of the significance of frailty in several aspects. We had an exceptionally long follow-up and observed that the frailty-mortality relationship extends up to 30 years from baseline. In addition, half our sample consisted of younger individuals who are understudied for frailty in comparison to old individuals. We demonstrated that FI was a strong predictor of mortality especially in younger women, where the risk conferred by the increase in frailty was greater than that among the older. This finding is in line with previous results (16) that demonstrated a similar phenomenon in an age, sex and education-adjusted model on 12-year mortality. Hence, our results further verify the usefulness of FI in early risk stratification. The reason why the association between FI and mortality was not statistically significant in older men is unknown, yet we identified a data-driven explanation for it. CVD was the major cause of mortality among older men and as the FI did not predict CVD deaths, the association with all-cause mortality was hence likely attenuated. Similar findings have been reported in a Finnish study where the association between the Fried frailty phenotype and 4-year mortality was no longer significant in old men after adjusting for covariates such as smoking, education and functional capacity (18). Nevertheless, it should be noted that both the FI and the frailty phenotype have been found to predict mortality in older men in several studies (1,2).

Our findings are in accordance with a recent meta-analysis reporting that both frailty and pre-frailty predict CVD-related mortality in individuals aged 65 years and older (9). Sex-differences were not, however, specifically addressed in that work. Related findings have been presented in patients with established CVD or after an acute cardiovascular event where the presence of frailty - defined using different indicators - has been an independent predictor of all-cause mortality (19,20). Although the individuals in these studies have generally been older than in our study and the follow-ups have been shorter, our findings are given some support by an Icelandic study that demonstrated a predictive validity of frailty on incident CVD, including deaths due to cardiovascular causes (10). The association was stronger in women than in men and independent of subclinical atherosclerotic manifestations. Because frailty and subclinical CVD may share common biological pathways, a causal relationship is unlikely. Nevertheless, a mutual reciprocal relationship has been proposed; frailty may lead to CVD and CVD can also lead to frailty (20). Notwithstanding that both scenarios are possible, our results provide new insight to the matter by demonstrating that FI predicts CVD mortality, regardless of CVD status at baseline, age and smoking. This observation may also be indicative of the type of vulnerability that the FI reflects in women. Given that our follow-up for cause-specific mortality spanned 27 years and the median time to CVD death in women was 14.6 years, our observation is of interest when considering FI as an early risk assessment tool.

The results on cancer mortality suggest an association with FI in women yet the matter warrants further confirmation as the number of cancer deaths in this sample was rather small. The association was also attenuated after adjusting for cancer diagnosis at study baseline, indicating that it was largely driven by those individuals who already had cancer. Small event frequency likely accounted for the non-significant association in the SHR model as the relatively higher FI levels among those experiencing death due to CVD and other causes (that were kept within the risk set) outbalanced the risk for cancer death. It is nevertheless noteworthy that despite the higher number of cancer deaths among men, no associations were observed. Our results in women align with the observations in a study of breast cancer patients where frailty was shown to be a predictor of both all-cause and breast cancer mortality irrespective of treatment differences between the robust and the frail (21). A systematic review has also demonstrated that frailty is an independent predictor of up to 10-year all-cause mortality in patients with various cancers (12). Together with these observations, our results give support to the notion that frailty can predispose to cancer-related vulnerability, possibly through treatment intolerance and postoperative complications (12).

The finding of a lower FI being predictive of dementia mortality in the SHR model may be attributable to the fact that those who died of dementia had longer median times to death compared to the other causes, which in the SHR approach translates to the risk set including an increased proportion of individuals who already died due to the other causes (and had higher FI levels). Those who died of dementia can also be considered as healthy agers as they escaped deaths due to other causes and survived to more advanced ages. Nevertheless, as the corresponding association in the CHR model was non-significant, our data do not allow conclusive inferences on this relationship. However, we can conversely argue that the FI does not capture the vulnerability to dementia death in community settings, yet in older ages frailty has been shown to be predictive of dementia diagnosis (11).

The only model where the FI conferred a similar increased risk in men and women was the CHR approach on other-cause mortality. In fact, the other-cause mortality was the only model where FI was significant in men. Moreover, the median FIs at baseline showed greater differences across the causes of death i.e., CVD, cancer, dementia and other, in women than in men. Hence, it appears that the FI-related mortality risk is distributed evenly between the different causes in men, whereas in women it is largely confined to CVD deaths. FI can thus be considered to represent a different type of vulnerability in men and women - a finding that may add to the understanding of the frailty paradox with regards to the greater overall risk of death in men across all levels of frailty (22).

In conclusion, our results demonstrate that that FI is a prognostic survivorship factor already at younger ages (<65 years) and that the association is stronger in women. When different causes of death are considered, FI can best identify women at risk of CVD death and to a lesser degree the risk of other-cause death. In men, FI can be understood as more of a general vulnerability factor as the risk appears to be evenly distributed between the causes and confined to the other-cause mortality. However, more research on this topic is required as our population was rather small for a cause-specific mortality analysis. Moreover, this was the first study to assess the predictive value of FI on mortality in a competing-risks setting.

## Acknowledgments

We thank Professor Kenneth Rockwood for his valuable input in creating the frailty index in our sample. We also thank Professor Chandra Reynolds and professor Deborah Finkel for careful reading of the manuscript and helpful comments.

